# The PREGCARE study: Personalized recurrence risk assessment following the birth of a child with a pathogenic *de novo* mutation

**DOI:** 10.1101/2022.07.26.501520

**Authors:** Marie Bernkopf, Ummi B. Abdullah, Stephen J. Bush, Katherine Wood, Sahar Ghaffari, Eleni Giannoulatou, Nils Koelling, Geoffrey J. Maher, Loïc M. Thibault, Jonathan Williams, Edward M. Blair, Fiona Blanco Kelly, Angela Bloss, Emma Burkitt-Wright, Natalie Canham, Alexander T. Deng, Abhijit Dixit, Jacqueline Eason, Frances Elmslie, Alice Gardham, Eleanor Hay, Muriel Holder, Tessa Homfray, Jane A. Hurst, Diana Johnson, Wendy D. Jones, Usha Kini, Emma Kivuva, Ajith Kumar, Melissa M. Lees, Harry G. Leitch, Jenny E. V. Morton, Andrea H. Németh, Shwetha Ramachandrappa, Katherine Saunders, Deborah J. Shears, Lucy Side, Miranda Splitt, Alison Stewart, Helen Stewart, Mohnish Suri, Penny Clouston, Robert W. Davies, Andrew O. M. Wilkie, Anne Goriely

**Affiliations:** Clinical Genetics Group, MRC Weatherall Institute of Molecular Medicine, Radcliffe Department of Medicine, University of Oxford; NIHR Oxford Biomedical Research Centre, Oxford, UK; Victor Chang Cardiac Research Institute, Darlinghurst, NSW, Australia; Centre for Population Genomics, Garvan Institute of Medical Research and School of Mathematics and Statistics, UNSW Sydney, Sydney, New South Wales; Centre for Population Genomics, Murdoch Children’s Research Institute, Melbourne, Victoria, Australia; Oxford Genetics Laboratories, Churchill Hospital, Oxford University Hospitals NHS Foundation Trust, Oxford, UK; Oxford Centre for Genomic Medicine, Nuffield Orthopaedic Centre, Oxford University Hospitals NHS Foundation Trust, Oxford, UK; Manchester Centre for Genomic Medicine, Manchester University NHS Foundation Trust and Division of Evolution and Genomic Sciences, University of Manchester, Manchester, Greater Manchester, UK; Department of Clinical Genetics, Liverpool Women’s NHS Foundation Trust, Liverpool, UK; Clinical Genetics Department, Guy’s Hospital, Guy’s & St Thomas’ NHS Foundation Trust, London, UK; Nottingham Regional Genetics Service, City Hospital Campus, Nottingham University Hospitals NHS Trust, Nottingham, UK; South West Thames Regional Genetics Service, St George’s University Hospitals NHS Foundation Trust, London, UK; North West Thames Regional Genetics Service, London North West University Healthcare NHS Trust, Northwick Park Hospital, Harrow, UK; North East Thames Regional Genetics Service, Great Ormond Street Hospital NHS Foundation Trust, London, UK; Sheffield Clinical Genetics Service, Sheffield Children’s NHS Foundation Trust, Sheffield, UK; Clinical Genetics, Royal Devon University Healthcare NHS Foundation Trust, Exeter, UK; MRC London Institute of Medical Sciences, Institute of Clinical Sciences, Faculty of Medicine, Imperial College London, London, UK; West Midlands Regional Clinical Genetics Service and Birmingham Health Partners, Birmingham Women’s and Children’s Hospitals NHS Foundation Trust, Birmingham, UK; Nuffield Department of Clinical Neurosciences, University of Oxford, UK; Wessex Clinical Genetics Service, University Hospital Southampton, Princess Anne Hospital, Southampton, UK; Northern Genetics Service, The Newcastle upon Tyne Hospitals NHS Foundation Trust, Newcastle, UK; Department of Statistics, University of Oxford, Oxford, UK

**Keywords:** mosaicism, gonadal mosaicism, de novo mutation, WGS, trio sequencing Personalised Recurrence Risk, Precision genetic counselling

## Abstract

Next-generation sequencing has led to a dramatic improvement in molecular diagnoses of serious pediatric disorders caused by apparently *de novo* mutations (DNMs); by contrast, clinicians’ ability to counsel the parents about the risk of recurrence in a future child has lagged behind. Owing to the possibility that one of the parents could be mosaic in their germline, a recurrence risk of 1-2% is frequently quoted, but for any specific couple, this figure is usually incorrect. We present a systematic approach to providing individualized recurrence risk stratification, by combining deep-sequencing of multiple tissues in the mother-father-child trio with haplotyping to determine the parental origin of the DNM. In the first 58 couples analysed (total of 59 DNMs in 49 different genes), the risk for 35 (59%) DNMs was decreased below 0.1% but for 6 (10%) couples it was increased owing to parental mosaicism - that could be quantified in semen (recurrence risks of 5.6-12.1%) for the paternal cases. Deep-sequencing of the DNM efficiently identifies couples at greatest risk for recurrence and may qualify them for additional reproductive technologies. Haplotyping can further reassure many other couples that their recurrence risk is very low, but its implementation is more technically challenging and will require better understanding of how couples respond to information that reduces their risks.

## Main text

The birth of a child with a serious clinical disorder to a healthy couple with no previous family history is a life-changing event. Added to the challenges posed by caring for their child, is the anxiety that their future children could be similarly affected. Whilst robust frameworks for addressing this possibility are increasingly available for common chromosomal abnormalities and recessive monogenic diseases, no systematic approach has been developed for dominant disorders caused by apparently *de novo* mutations (DNMs). Such disorders are collectively common, estimated to affect at least 1 in 295 births^1^, but extremely heterogeneous; for example, mutations in over 650 genes are currently recognised to cause developmental disorders through a dominant mechanism of action^1,2^. The need to address this issue has been made more pressing by the success over the past decade of next-generation sequencing (NGS) technologies in identifying DNMs, leading to a deluge of new causative genes and diagnoses.

The implementation of NGS technologies across large populations has contributed to a better understanding of the patterns of occurrence of DNMs. It is now well established that DNMs are rare events (spontaneous human mutation rate is ∼1.2 × 10^−8^ per bp, per generation), mainly occurring as “one-off” copying errors during sperm production, or less frequently in oocytes^3,4^. While in these instances, the risk of recurrence for future siblings would be negligible, DNMs can also occur post-zygotically (either in one of the two clinically unaffected parents, or in the affected child) leading to a mosaic genotype that alters the recurrence risk. Mosaicism populating multiple germinal cells in the ovaries or testes (arising during one of the parent’s own development), termed gonadal (or germline) mosaicism, may be associated with a substantial recurrence risk for further offspring, reaching up to 50% in some cases; by contrast, convincing demonstration of post-zygotic mosaicism in the offspring would eliminate the chance of sibling recurrence^5^.

Although mosaicism has long been recognised as a source of DNMs, few studies have attempted (or had the power) to define its exact contribution to spontaneous disease. Overall, current NGS methods used to identify DNMs rely on mother-father-proband trio sequencing and are poorly suited for detection of mosaic cases - either for cases of low-level (parental) mosaics^6^, or to distinguish high-level variant allele frequency (VAF) from constitutional (50%) presentation in post-zygotic (proband) cases^5,7^. For example, the limit of VAF sensitivity of WES/WGS trio sequencing, which is typically performed at a depth of 25-30x, is ∼10-15%, similar to that of dideoxy-sequencing^6,8^. Moreover, routine genetic analysis relies on the interrogation of a single somatic tissue (blood or saliva), which is not adequate to identify mosaicism in parental gametes or variable VAF in a proband’s tissues.

The recognition that the tissue distribution and VAF of a DNM are determined by the timing at which it first occurred, allows us to identify three key time points during development with different predicted presentations: (1) very early in development - before the segregation of germline and somatic lineages at ∼day 14 of human embryogenesis, yielding cases of mixed (somatic and gonadal) mosaicism; (2) post-15 days of development in the germline, resulting in confined gonadal mosaicism; (3) or much later in the developing or adult gonad, yielding a “one-off” mutation (Supplementary Fig S1). Furthermore, by taking into account the individual in whom the DNM originated (mother, father, or affected child), it becomes possible to distinguish a total of seven scenarios whereby a DNM can occur (Fig. 1). The overall relative prevalence of these seven scenarios can be estimated quite accurately based on previous analyses of the parental origin of DNMs and the prevalence of mosaicism from population studies (see Supplementary Note 1).

**Fig. 1:**
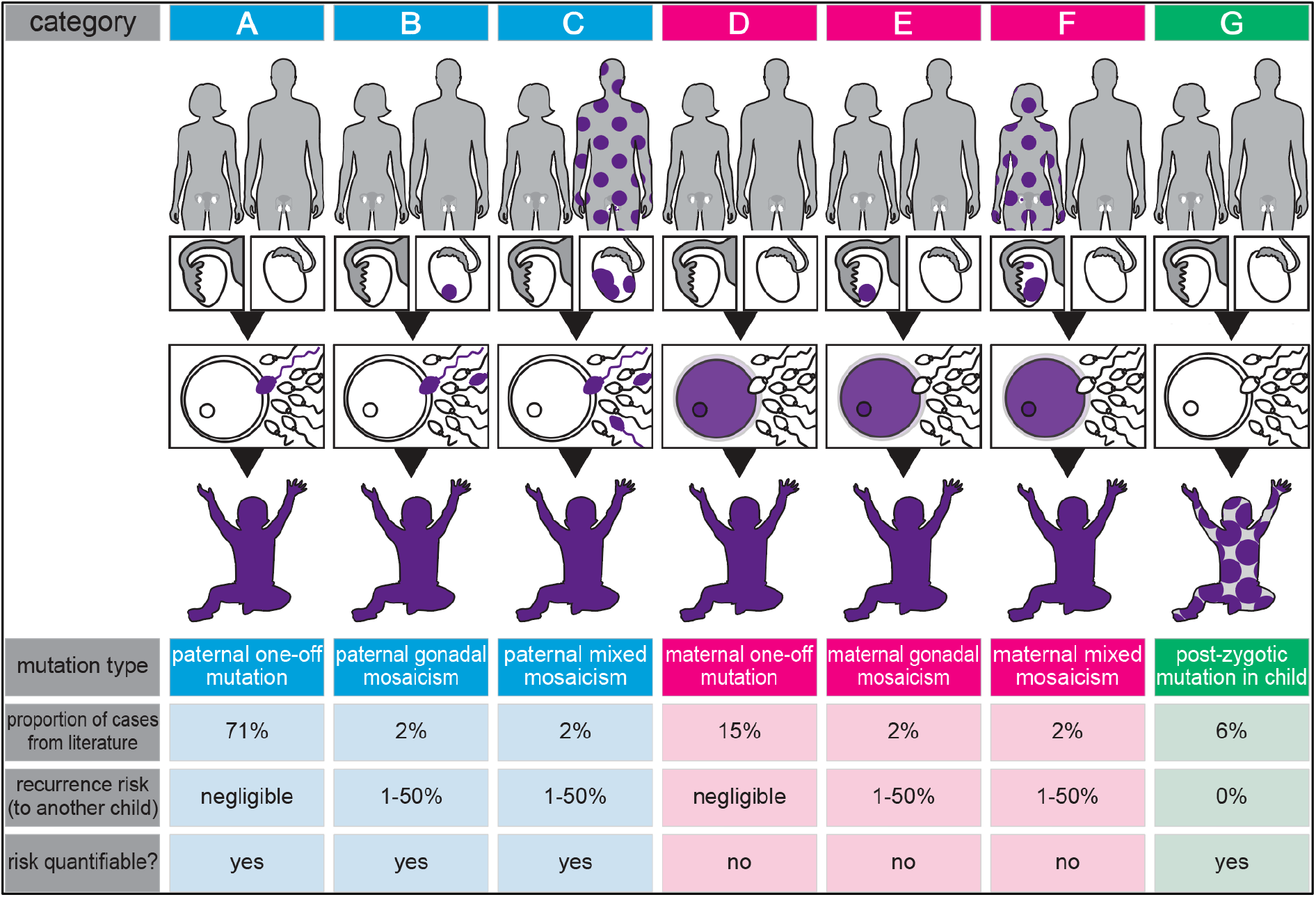
Stratification of DNMs into seven categories. Establishing the origin (paternal (blue), maternal (pink) or post-zygotic (proband, green), and timing of the mutational events (purple colour indicates mutant cells), yields widely different recurrence risks in different families. See main text, Supplementary Fig S1and Supplementary Note 1.

Here, we have developed a systematic strategy to categorise pathogenic DNMs in a mixed clinical population of 60 couples who had one or more children with a serious developmental disorder caused by an apparent DNM, and were seeking individualized reproductive counselling about recurrence risk in a future pregnancy. By combining deep-sequencing of multiple tissues to detect occult mosaicism with haplotyping to determine parent-of-origin of the DNM, we show that we can reliably stratify individual couples into discrete categories that are associated with substantially different risks to the offspring. This personalised approach to recurrence risk assessment offered *prior* to a new pregnancy should provide reassurance to the majority of couples in whom the risk is very low or negligible and help to focus resources on the minority of families at increased recurrence risk.

## Results

### Population sampled

Following ethical approval we recruited, through the network of Clinical Genetics centres in England, 60 couples who had one or more children (or fetuses) affected by a serious clinical disorder caused by an identified DNM, which was not present in the parents’ DNA on routine analysis (PREGCARE study; Online Methods). Two families (FAM17 and FAM60) had three affected siblings/pregnancies, indicating that one of the parents must be a gonadal mosaic, but routine diagnostic analysis performed on parental blood DNA had failed to identify the parent-of-origin. To eliminate ascertainment bias, these two families are excluded from the quantitative presentation of the data but included in the specific analysis of mosaicism. Hence, our primary cohort comprises data from 58 parent-child trios, including one trio with two different pathogenic DNMs (FAM12). These 59 DNMs comprised 40 single nucleotide substitutions, 14 small (1-2 nucleotides) indels and 5 larger (4-44 nucleotides) indels in 49 different genes, providing a broad and representative spectrum of pathogenic molecular lesions encountered in clinical practice (Supplementary Table S1).

### Deep-sequencing of multiple tissues identifies mosaic cases

Four of the seven categories shown in Fig. 1 (B, C, F and G) involve mosaic states that can be directly identified without requiring invasive sample collection and distinguished by deep-NGS of tissues collected from the family trio. We therefore obtained up to 14 biological family samples (child: blood, buccal mucosa left + right; mother and father: blood, saliva, buccal mucosa left + right, urine; plus paternal semen) to seek evidence of mosaicism. Collection of parent samples was designed to include all three embryonic germ layers (ectoderm, buccal; mesoderm, blood; endoderm, urine), plus germline in the father (Supplementary Fig. S1).

The overall strategy deployed for the analysis is shown in Fig. 2.

**Fig. 2:**
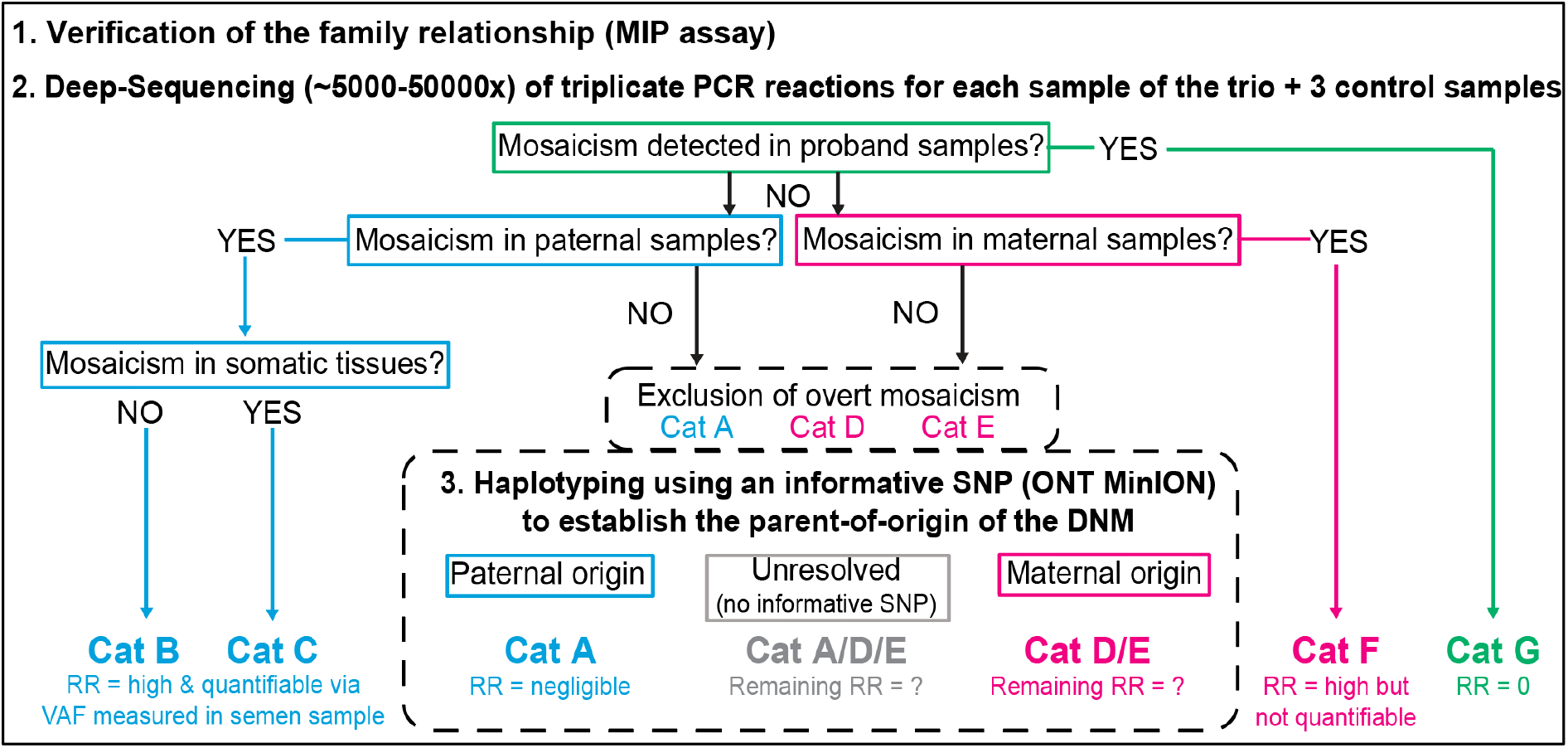
Flow chart describing the three-tier sample analysis in the PREGCARE study. Following collection of up to 14 different biological samples per family and verification of the familial relationships between the 3 individuals of the trio, the DNM site was deep-sequenced in all family samples (performed in triplicates together with 3 unrelated controls) to detect low-levels of parental mosaicism or instances of post-zygotic mosaicism in the proband. For those families without evidence of overt mosaicism, haplotyping was performed to resolve the parental origin of the DNM and further stratify the recurrence risk (RR). Refer to Fig. 1 for category (Cat) description.

Following verification of sample relationships and parentage in each family using a panel of bespoke multiplex inversion probes (MIP) targeting 168 common single nucleotide polymorphisms (SNPs), we designed a custom PCR assay covering 65-224 bp around the family-specific DNM site and performed triplicate reactions from each available tissue, and three unrelated control DNAs, before undertaking deep-NGS (target depth 5,000-50,000x) on the Illumina MiSeq platform. Reads were processed using amplimap^9^ and VAF quantified at the genomic position of the DNM (see Methods). NGS was poorly suited to analyse two DNMs associated with the larger indels (a 35 bp deletion in FAM12b and a 44 bp duplication in FAM54). Hence, to rule out the possibility of occult mosaicism in these samples, we performed mutant allele-specific PCR on all available samples from the trio (Supplementary note 2).

Overall, deep-NGS (and/or allele-specific PCR) identified 7/59 (11.9%) cases with strong evidence of mosaicism in one family member (Fig. 3; Supplementary Fig. S2 & Table S2). These comprised DNMs belonging to Categories B (paternal gonadal mosaicism; FAM27), C (paternal mixed mosaicism; FAM34, FAM49, FAM58), F (maternal mixed mosaicism; FAM01, FAM50) and G (post-zygotic mosaicism in proband; FAM33). Analysis of the two additional families in which recurrence in siblings was already documented (FAM17, FAM60) showed that both were attributable to maternal mixed mosaicism (Fig. 3).

**Fig. 3:**
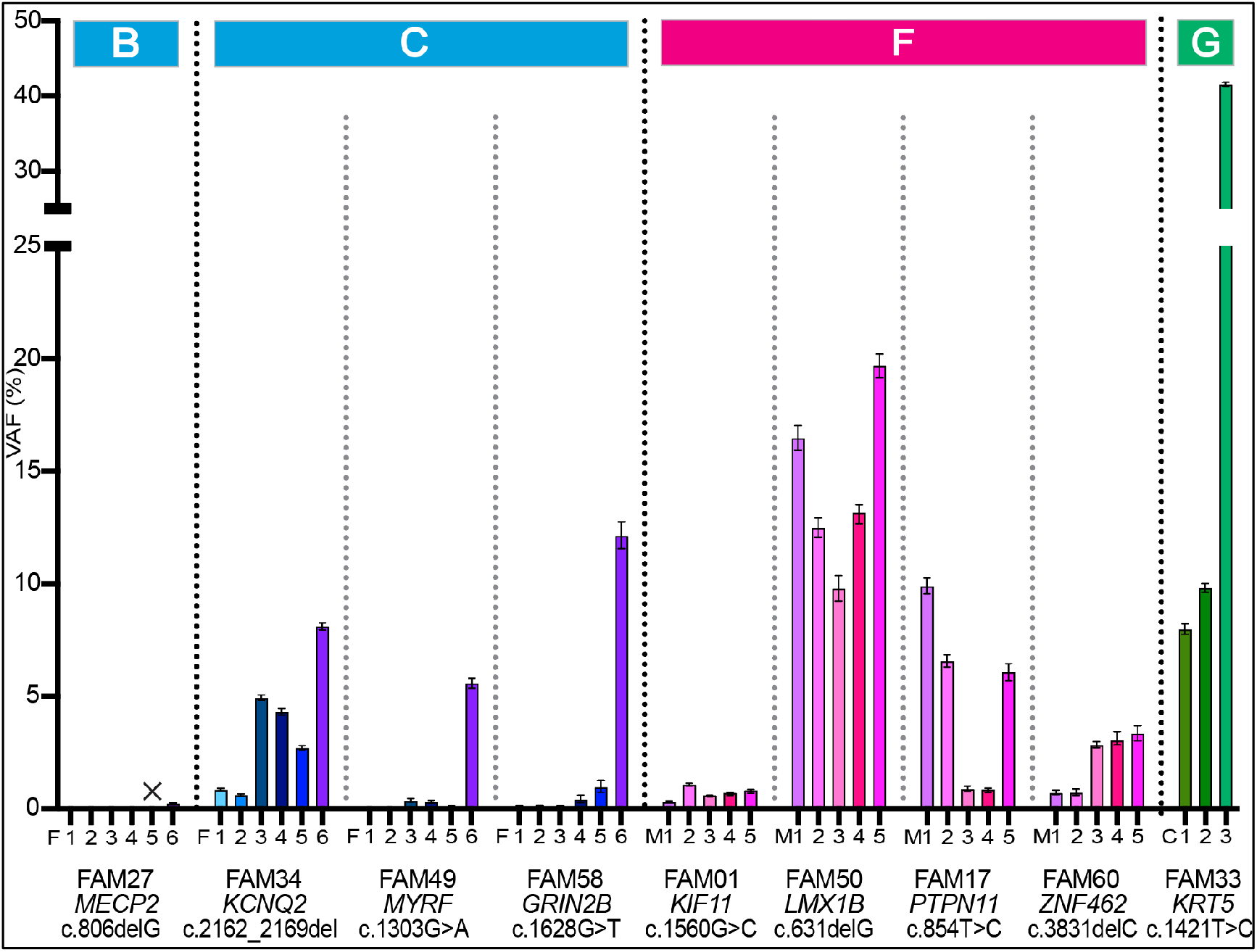
Mutation levels observed in the families presenting with mosaicism. Variant allele frequencies (VAF) in different samples from the family member in whom mosaicism was detected by deep-NGS sequencing. Family number, gene, cDNA coordinates of the DNM and the origin of the different samples are indicated on the x-axis in the same order for each family and distinguished by colors for ease of visualization. The category classification is indicated at the top of the figure. X represents a sample failure. Full data for the other family members and controls are presented in Supplementary Fig S2 and Table S2. FAM17 and FAM60 are the two families with multiple affected pregnancies and belong to category F (maternal mixed mosaicism); note the low VAF in the maternal blood samples (M3) for both families. The corrected VAF (see Methods) is plotted and error bars represent the 95% binomial confidence intervals across the three technical replicates. Abbreviations: F=father; M=mother; C=child; 1=buccal swab (left); 2=buccal swab (right); 3=blood; 4=saliva; 5=urine; 6=sperm; 7=gDNA from original testing.

Identifying these mosaic families is particularly important, because whereas the recurrence risk associated with post-zygotic mosaicism (Category G) is effectively zero, the other three mosaic categories (B, C, F) are associated with increased recurrence risks. While the offspring risk is not directly quantifiable for the maternal mosaics because of the inaccessibility of ovarian tissue, it could be quantified in the paternal mosaic cases via the VAF measured in sperm and ranged from 0.23% (FAM27) to 12.1% (FAM58) (F6 bars in Fig. 3; Sup Fig S2). Importantly, in the three paternal cases of mixed mosaicism the level of the cognate DNM in sperm (5.6-12.1%) was substantially higher than in any of the other tissues sampled and variability in mutation levels was present between different somatic tissues, with no one tissue providing a reliable indicator of the level in sperm (Fig. 3). In seven of the eight parental mosaic cases, where DNA derived from blood (the most widely used source of DNA for genetic analysis) was analysed, the level of mutation in the transmitting parent was below 5% (F3 and M3 values in Fig. 3; Supplementary Table S2). Such VAFs would be impossible to detect systematically using standard diagnostic NGS read depths (∼25-30x) or dideoxy-sequencing, illustrating the importance of deep-sequencing (>5000x) and the value of collecting additional tissue samples to increase sensitivity for ascertaining occult mosaicism. In the single identified instance of paternal confined gonadal mosaicism, a relatively low level of sperm mutation was observed (0.23%), consistent with a slightly later timing of mutational origin (Supplementary Fig S1) and in line with empiric data on mutations in sperm^10-13^.

Also of note, the VAFs for the proband samples from FAM33 with the post-zygotic mutation (C1, C2 and C7 values in Fig. 3) were markedly different across the tissues analysed (blood 41.6%; buccal mucosa 8.0% and 9.8%), demonstrating the benefit of analysing several tissue samples from an individual to distinguish post-zygotic mosaicism associated with high VAF levels from constitutional (50%) presentation.

### Haplotype phasing enables determination of parental origin of DNM

For the remaining 52 DNMs that did not classify into one of the four mosaic categories described above, further stratification was attempted through haplotyping to determine the parental origin of the mutation (Fig. 2). For these families, only one category (Category E, maternal gonadal mosaicism) is associated with a recurrence risk to offspring (Fig. 1). Although it is not possible to distinguish Categories D and E (because oocytes are not accessible), most of the remaining DNMs (∼88%) are predicted to belong to Category A (paternal one-off), which is associated with a negligible risk to offspring.

Parental origin could be inferred for two families without performing haplotyping: FAM26 (mutation in the X-linked *MID1* gene in a male proband, implying a maternal origin) and FAM54 (a 34 bp duplication in *MAGEL2*, a gene known to be maternally imprinted and for which pathogenic mutations are exclusively paternal in origin^14^).

To perform haplotyping of the other 50 DNMs, we sought an informative SNP or other variant in close proximity to the DNM, to enable phasing of the parental alleles. In the most common informative scenario, the child is heterozygous for the SNP (genotype AB) whereas one of the parents is homozygous (genotype AA or BB), making it possible for the inherited parental chromosomes to be distinguished. In three cases an informative SNP was present in the PCR product used for the deep-sequencing, enabling the parental origin to be determined directly by examining the phase of the DNM on the Illumina reads. To haplotype the 47 remaining DNMs, we designed two long PCR products extending away on either side of the DNM (total genomic region covered ∼7-30kb), and sequenced the resulting fragments for the three family members using the MinION platform from Oxford Nanopore Technology (ONT). Reads for each trio were processed and analysed with an in-house custom pipeline combining Medaka and pile-up processing (see Methods and Supplementary Note 3). This haplotyping strategy was successful in the majority (38/47) of cases (Supplementary Table S3), including three families (FAM11, FAM38, FAM67) that required a more complex analysis involving two SNPs to distinguish the parental alleles. In one of these (FAM38), due to the local genomic context of the DNM (a single G-nucleotide deletion within a homopolymeric region), phasing by direct analysis of sequencing traces could not be resolved by ONT sequencing. Nevertheless, this approach identified an informative SNP in the proband which was used to design a bespoke allele-specific PCR and determine the DNM parental origin (Supplementary Note 4 & Table S3B).

Overall, parental origin could be established for 82.7% of DNM (43/52), which included 34 DNMs of paternal origin (79%) and 9 (21%) present on the maternally-derived allele (Fig. 4) - a result in line with the expected ∼4:1 male to female ratio of mutational origin^3^ (Supplementary Note 1).

**Fig. 4:**
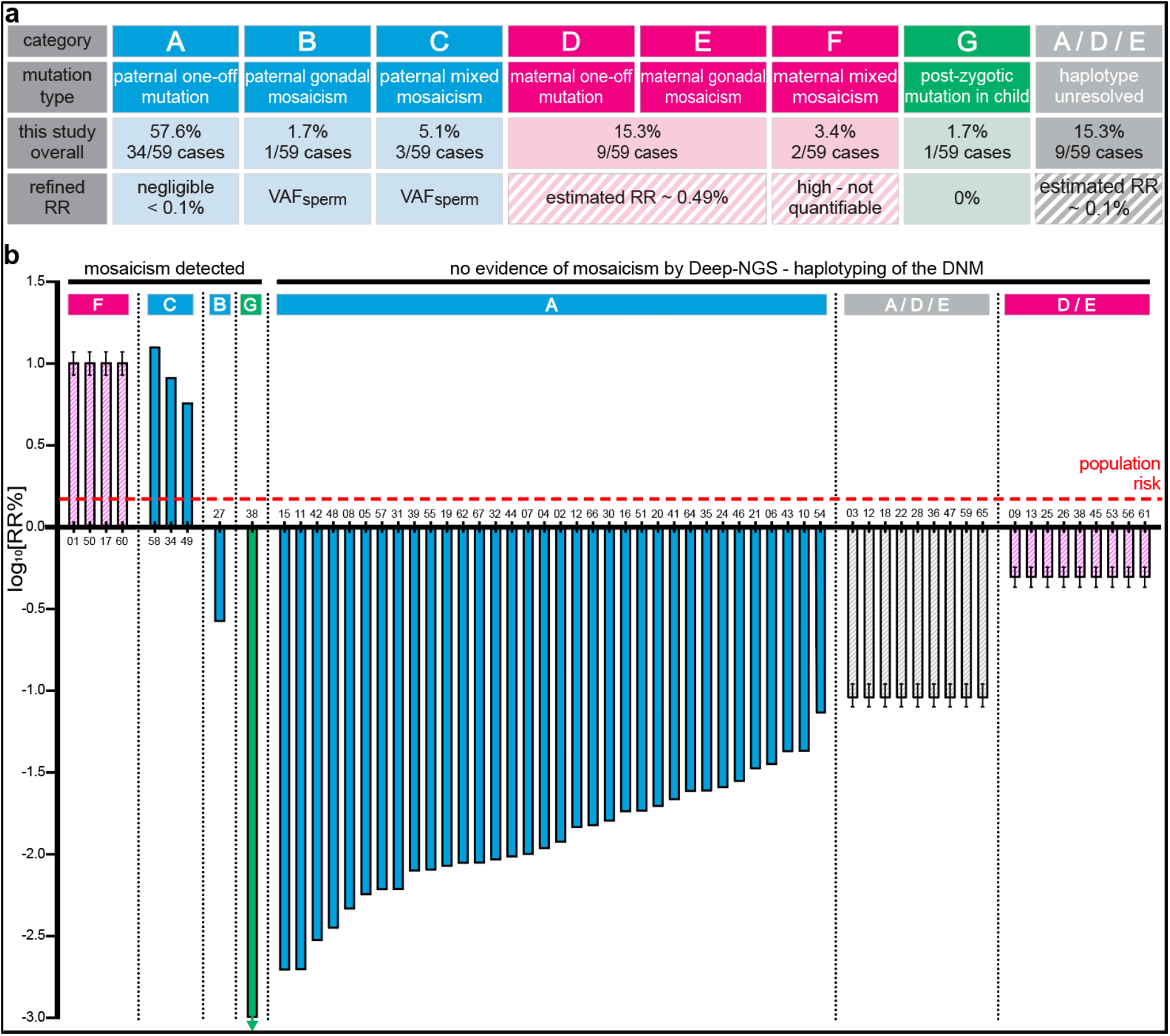
Overview of the results of the PREGCARE study showing refinement of individual recurrence risk for all families. **(a)** Summary table of the PREGCARE results for the 59 DNMs analyzed in this study and overview of the refined recurrence risk (RR). **(b)** Personalized recurrence risk (RR%) estimates for each of the 60 families (61 DNMs) enrolled in the PREGCARE study represented on a logarithmic scale. The red dotted line represents the generic population RR given to couples who have had a child with a DNM (∼1.5%). The RR can be quantified (block colours) for paternal (Categories A-C, via semen analysis) and post-zygotic (G) DNMs; note that to be conservative in our estimates of RR, we have plotted the upper 95% binomial CI from the corrected VAF_Sperm_ measured by deep-NGS for the paternally-derived DNMs (Supplementary Table S2). The RR can only be estimated for maternal (pink) or haplotype-unresolved cases (grey). These estimates are represented by stripes, with error bars representing the upper and lower 95%CI - see Supplementary Note 5. Note that the DNM of FAM54 was analyzed by allele-specific PCR (Supplementary Note 2) and that Category F includes the two additional families with multiple affected pregnancies. Individual family numbers are indicated on the x-axis.

### Combining deep-sequencing and haplotyping allows category stratification and individualized recurrence risk

Having singled out the mosaic cases by deep-NGS of multiple familial tissues, the particular value of the combined approach of deep-sequencing of semen samples with haplotyping is to identify those families in which the DNM is paternal in origin (34/52), as they belong to Category A (Figs. 1 and 2). Deep-sequencing of sperm of these paternal mutations allows measurement of the VAF and derivation of the upper confidence limit for the level of DNM present in sperm (Supplementary Table S2). As NGS is subject to background sequencing errors, we corrected the raw VAF values using measurements from three unrelated control samples. The corrected VAFs were estimated by numerically maximizing their marginal likelihood and 95% confidence intervals were obtained by using profile likelihood (see Methods for details). For Category A samples, the upper bound (95% CI) of the VAF measured by deep-NGS in sperm was below 0.05% in all cases (Supplementary Table S2; Fig. 4). These data point to DNMs in this category having originated as ‘one-off’ events during late gonadal development or adult spermatogenesis.

In 9/52 (17.3%) families, the haplotyping revealed a maternal origin of the DNM. In these cases, the negative findings from deep-NGS of maternal somatic tissues (i.e. maternal origin of mutation but with no evidence of somatic mosaicism) do not allow categories D and E to be distinguished (Figs. 1 and 2; 4). The relative risk for a DNM belonging to Category E (maternal gonadal mosaicism) and representing a recurrence risk in a future pregnancy, rather than to Category D (‘one-off’ maternal event) can be estimated to occur in ∼2:15 families (Fig. 1 and Supplementary Notes 1 & 5), meaning that on average 1 in every 8 or 9 maternal DNMs is anticipated to have originated in early developing germ cells. While the prevalence of mosaicism has not been directly quantified in ovaries owing to their experimental inaccessibility, it can be assumed that these early events are similar in magnitude for ovaries and testes because germline lineages are specified several weeks prior to sex determination.^4^ To gain further insight into the remaining recurrence risk for DNMs of proven maternal origin (D/E), we used estimates of VAFs observed for paternal confined mosaics obtained from sperm WGS^11^ (Supplementary Note 5). We obtained a recurrence risk estimate for the combined maternal categories D-E of 0.49% (95%CI: 0.43% - 0.57%), a modest reduction over the population average (Fig. 1 and Fig. 4).

### Evaluation of the remaining risk for cases with unresolved parental origin

Finally, in a further 9 families, despite sequencing a ∼13-27 (mean 22.2) kb region around the DNM in the proband, no informative SNP could be identified in the MinION reads. Hence the parental origin of the DNM could not be assigned, and the mutation could belong to any of Categories A, D or E (Fig. 1). As the majority of DNMs are predicted to be sporadic (Categories A or D), the remaining risk (associated with Category E) for these couples can be estimated by combining the relative proportion of Category E cases (2:86) and the average VAF observed for gonadal mosaicism^11^. As a result, for a DNM with an unresolved parent of origin, the recurrence risk is estimated to be 0.09% (95%CI: 0.08%-0.11%), a reduction of approximately 10-fold compared to the population risk baseline (Fig. 4; Supplementary Note 5).

## Discussion

We have applied a general framework to analyse systematically and at scale, the origins of DNMs presenting in a clinical setting. The work addresses a stark unmet clinical need to improve genetic counselling for couples who have had a child affected by a disorder caused by a DNM – a situation faced by almost a million parents annually – in order to provide them with a personalised risk assessment *prior* to a new pregnancy. The current standard of care, which is to provide these couples with a recurrence risk of ∼1-2% is unsatisfactory, both because this figure is nearly always wrong (as illustrated by Fig. 4), but also because of the uncertainty it raises for the complex decision process of whether to extend their family. It is well documented that couples’ attitudes to reproductive risk vary widely^15^: some will view the 1-2% risk as small and others would not contemplate extending their family in the face of any risk. In addition, while in many healthcare settings there may be the option of a prenatal diagnostic procedure (chorionic villus biopsy or amniocentesis), this is associated with a small risk of miscarriage (currently estimated as ∼0.2-0.5% for each procedure^16-18^) and may not be ethically acceptable to some couples. Owing to a combination of cost and technical challenges, prenatal procedures that are non-invasive (assay of free fetal DNA from maternal blood sample) or those which avoid the possibility of termination of pregnancy in the event of recurrence (preimplantation genetic testing for monogenic disorders [PGT-M]), are not available in most public healthcare settings. For example, in the UK the eligibility threshold for PGT-M is a risk >10% of having a child with a serious genetic condition, which excludes the parents of children with DNMs even though some couples will have a risk higher than this.

Over recent years several pioneering studies on DNM origins have provided a solid framework to quantify the relative contribution of different mutational processes to DNMs (Fig. 1; Supplementary Notes 1 & 5)^3-5,19-21^. We designed the PREGCARE study based on this framework, with the dual aims to seek evidence for mosaicism in each member of the parent-child trio (deep-sequencing), and to stratify the risk based on the likely timing and parental origin of the DNM. Important aspects of the study design include the recognition that (1) clinically-relevant mosaicism is caused by early embryonic mutations, that present either in both soma and germline (mixed mosaicism) or the germline only and affect males and females equally - because they originate before sex determination; (2) sampling of multiple tissues of different embryonic origins increase the likelihood of detecting instances of mixed mosaicism in parents (or post-zygotic events in the proband); and (3) analysis of a paternal semen sample allows direct quantification of risk for paternally-derived DNMs, which are anticipated to represent ∼3/4 of cases. Hence, although the female germline is not accessible to direct analysis, data about the prevalence and VAF anticipated for maternal mosaic cases can be inferred from sperm data^11^. Moreover, the relative risks of mixed vs. confined gonadal mosaic events can be estimated based on data from deep-sequencing of paired blood and sperm samples,^10,11^ which have shown that the average VAF measured in sperm for cases associated with mixed mosaicism is ∼9%, while ‘sperm-only’ average VAF are ∼3%^11^ (Supplementary Note 5). These data suggest that because mixed mosaicism is caused by very early mosaic events, they are likely to have a wide tissue distribution and be present at higher VAF (Supplementary Fig S1). Moreover, the rate of spontaneous mutations may be elevated during the first embryonic divisions^22,23^. Hence, mixed mosaic cases likely contribute to most of the recurrence risk in the next generation and identifying them by deep-sequencing of somatic tissues (and semen) represents an efficient way to single out the couples at higher risk.

Here we show through systematic analysis of a clinical series of 59 DNMs a very good correspondence between the distribution of DNMs across the 7 different categories for the families analysed to that anticipated from previous work (Figs. 1 & 4A; Supplementary Note 1). In our cohort, which consists of clinically-ascertained cases, DNMs originated from occult parental mosaicism in ∼10% (6/59) of cases. For five families it was detectable in the transmitting parent’s somatic tissues - although present at low VAFs in blood, illustrating the importance of deep-sequencing (>5000x) and the value of collecting additional tissue samples to increase sensitivity for ascertaining occult mosaicism.

Fig. 4, which summarises our overall findings, shows that we achieved risk alteration for individual couples over more than three log10 orders of magnitude: for 54/59 DNMs, the risk was reduced compared to the population baseline risk, and for 5/59 (the mixed mosaics), it was likely increased (but only quantifiable in the 3/5 paternal cases).

Encouraging though these data are, we acknowledge several barriers before considering clinical translation of this work. The first hurdle relates to technical implementation of individualised recurrence risk measurement in a clinical setting, which requires robust laboratory methods and will be challenging as a DNM-specific custom assay will be required for most families. In this study we used two methods, deep-sequencing and haplotyping, that provide complementary information. Deep-NGS is highly effective in singling out couples at high recurrence risk, whereas haplotyping is essential to generate most of the very low recurrence risks and reassure the majority of couples that they belong to Category A. The Illumina platform used for deep-sequencing is technically straightforward and the associated calling pipelines are readily available in most diagnostic settings. Of note, for efficient evaluation of reproductive risk, the source of tissue samples ought to be a major consideration. While semen provides the ideal tissue for determining reproductive risks directly, surprisingly at present the prevailing working practices of clinical genetics do not include routine semen analysis. Our view is that this work and that of others^11-13,24^ provides clear evidence to promote much more widespread collection and analysis of this material (as is standard, for example, in fertility clinics). As there is no easy access to maternal gonadal tissue, it would be valuable to know whether there is a particular somatic tissue that provides a better surrogate for the germline. This can be addressed by studies of male samples, but although we observed substantial variation, it was not possible to identify a clear surrogate^12,13,25^, consistent with the fact that during early embryogenesis, cell populations are subject to bottlenecks and differential lineage commitments leading to considerable variation and stochasticity in cellular representation across tissues^20,26^. Hence reliance on assessment of a single tissue (blood) risks missing some mixed mosaics harbouring low mutation levels (or high level post-zygotic mosaicism in the proband). Of the other tissues we sampled, we found that saliva tended to reflect the results from blood^6,27^ but occasionally exhibited a higher background that can bias low VAF interpretation, likely reflecting the fact that ∼70% of saliva DNA is derived from white blood cells, while the remaining fraction contains bacterial and/or other genomes (potentially including that of other family members, including the proband)^28^. Unlike urine, which often yielded poor amounts of DNA, buccal brushings (left and right sides of the cheek sampled independently), are easy to collect (including from children), and store, and contained cells of a different embryological origin to blood, which often yielded informative data.

Overall, we conclude that clinical implementation of deep-sequencing of a few key tissues from the trio (blood, buccal brushings and paternal sperm) alone should be easy to achieve and would identify most of the high-risk cases, therefore reducing the risk of mosaicism presentation for the remaining couples (Categories A, D, and E) to ∼0.1% (Supplementary Note 5).

To further refine the remaining risk in non-mosaic families, we used the ONT platform as a second method in this study. Although harder to scale and process than Illumina data, ONT showed good potential for implementation in diagnostics, but is not currently approved for use in most clinical settings. Independently of technical considerations, one major limitation of this approach, which led to a substantial minority (15.3%) of unresolved cases in this study, is the requirement for the presence of a heterozygous SNP in the vicinity of the DNM to distinguish the two parental alleles in the proband. Implementation of novel ultra-long read WGS methods will facilitate SNP identification and systematic parent-of-origin assignment of DNMs, but are not currently available in most settings^29^.

Another potential barrier to clinical implementation relates to how these refined risks are viewed by couples and whether changes in risk actually result in altered decision-making. Concerning the accuracy of our risk estimations, among the 61 DNMs analysed, in only 39 do we consider the risks to be reasonably accurate; these include the 38 DNMs shown to be paternally originating, in which we could directly measure levels of mutation in sperm, and the single post-zygotic case (Fig. 4). In 36/39 we reported a risk lower than baseline, while in the three mixed mosaic cases it was increased (to 5.6%, 8.1% and 12.1%). By contrast, in the remaining cases shown either to be of maternal origin (13/61) or unresolved (9/61), the risk estimation has been refined but remains inaccurate and may be viewed differently by parents and healthcare professionals. Even in proven cases of maternal mixed mosaicism, VAFs in somatic tissues are poor predictors for the germline, as illustrated by the two families with multiple recurrences in whom we detected relatively low VAF in maternal somatic tissues (maximum of 3.3% and 9.9% in the samples analyzed), despite three affected pregnancies in each sibship (Fig. 3). Nevertheless, detection of mixed mosaicism in maternal tissue will warrant caution in future pregnancy and it should also be noted that some diagnostic options may be more complicated for these families because of the unsuitability of non-invasive prenatal testing via analysis of cell-free fetal DNA in maternal plasma^30^.

In those cases where somatic mosaicism has been excluded but the DNM is proven or possibly maternal in origin, the risk of maternal gonadal mosaicism (Category E, Fig. 1) may remain an important factor in decision-making, despite the relative reduction in risk for these subcategories (Category E corresponds to ∼2:15 maternally-proven DNM and ∼2:88 haplotype-unresolved DNM with an estimated average VAF of ∼4%).

An interesting illustration from this work of the complexity of recurrence risk counselling is provided by the case of paternal confined gonadal mosaicism (Category B) in FAM27, in which the risk for the *MECP2* mutation was found to be 0.23% (95% CI: 0.19%-0.26%), over 5-fold lower than the 1.2% baseline population risk. How to counsel a couple in this situation, where stratification to the “at risk” Category B predicts increased caution, remains difficult. This risk may also need to be leveraged against the current UK recommendation for ‘higher risk’ in respect to the probability of carrying a fetus with Down Syndrome for a 35-year old mother ∼ 1/150 (0.66%) for which prenatal screening is routinely offered^31^.

Overall, we show that providing pre-conception recurrence risk assessment to couples who have had a child with a DNM can be achieved and offers the prospect of driving a major transformation in the practice of genetic counselling. Our data demonstrate that for all couples, it is possible to refine the risk of having another affected child with the same DNM and in the majority of cases (>64%) the risk is in fact very small, potentially reducing anxiety and the need for expensive pre-implantation or prenatal diagnostic options. For couples in whom we detected overt mosaicism, the risk is higher (and quantifiable through sperm analysis for the paternal cases). Providing evidence-based estimation of the actual risk will allow these couples to be singled out for further investigations and support, allowing them to make informed choices (and for their clinicians to provide them with personalised advice and risk assessment) about the different diagnostic options available to them.

## Acknowledgments

This work was primarily supported by grants from NewLife (17/18/04), the Wellcome (102731/Z/13/Z and 219476/Z/19/Z), and the National Institute for Health Research (NIHR) Oxford Biomedical Research Centre Programme. We acknowledge WIMM core funding from the Medical Research Council (MRC) through the WIMM Strategic Alliance (G0902418 and MC_UU_12025) and the support of the NIHR UK Rare Genetic Disease Research Consortium (Musketeers Memorandum). We thank Tim Rostron and the members of the Oxford Genomic Medicine laboratory for technical support.

The funders had no role in study design, data collection and analysis, decision to publish, or preparation of the manuscript.

## Authors contribution

Conceived the study and designed the experiments: AOMW and AGo

Performed the experiments: MB, UBA, KW and SG

Performed data analysis: MB, UBA, SJB, EG, NK, GJM, LMT, JW, RWD, AOMW and AGo

Recruited participants and/or provided samples: JW, EMB, AB, EB-W, FBK, NC, ATD, AD, JE, FE, AGa, EH, MH, TH, JAH, DJ, WJ, UK, EK, AK, MML, HGL, JEVM, AHN, SR, KS, DJS, LS, MS, AS, HS, MSu, PC

Wrote the manuscript: AOMW and AGo with input of MB, SJB, RWD

## PREGCARE: Online Methods

### Recruitment into the PREGCARE study

The PREGCARE (PREcision Genetic Counselling And REproduction) study was approved by the London - Queen Square Research Ethics Committee under the reference number 17/LO/1025 (IRAS reference: 225264). Couples with one (or multiple) children, stillbirths or terminated pregnancies affected by a likely pathogenic *de novo* mutation (DNM) and who were potentially interested in personalized transmission risk assessment for future pregnancies were invited to participate by healthcare professionals during routine clinical genetic consultation. A DNM was defined as a single-nucleotide or small insertion-deletion variant detected in the proband that was absent in the parents’ DNA on routine diagnostic genetic analysis. DNMs occurring in one of six paternal age-effect genes (*FGFR2, FGFR3, HRAS, KRAS, PTPN11, RET*) were excluded, unless there were multiple affected pregnancies^1^. Couples where the mother was pregnant at the time of sample collection, those who were not both the biological parents of the affected child, or either the biological mother or father did not consent to participate, were also excluded.

Recruitment and sample collection took place at 13 of the 17 participating National Health Service (NHS) Trusts in England, UK.

### Sample collection

Families interested in participating in the study were sent a box containing kits and instructions for collection at home of 2 ml saliva (Oragene DNA, OG-500, DNA-Genotek, Canada) and 50 ml morning midstream urine (Urine Collection And Preservation Tube, Norgen Biotek Corp., Canada) from both the mother and father, and an ejaculate of semen (following abstinence for three days before collection and stored at -20 °C) from the father. During the clinical visit, informed written consents were obtained and further samples were collected from the three family members, including 5 ml peripheral blood (EDTA) from father and mother and buccal cells from the left and right inner cheek lining from mother, father, and the affected child using swabs (sterile PurFlock Ultra tip swab in dry transport tube, Puritan Medical Products, ME, USA). Samples and completed consent forms were sent at room temperature to the MRC Weatherall Institute of Molecular Medicine where they were witnessed-transferred and processed for extraction or long-term storage within 48 hours of collection. Overall, a total of 67 boxes were dispatched and 60 families completed collection and consents and were enrolled into the study.

In addition, the child’s genomic DNA originally used for the molecular diagnosis was requested from the NHS genetic laboratory. This sample had usually been extracted from the proband blood or, occasionally, fetal tissues, amniocentesis or chorionic villus sample (CVS) (for details see Supplementary Table S1).

### Sample processing and DNA extraction

Upon delivery of the box to the lab, the family samples were given a unique identifier and processed. The saliva samples were incubated at 50 °C for 60 min and then aliquoted. Blood samples were aliquoted as whole blood and isolated buffy coat. Urine samples were centrifuged at 2000 x *g* for 10 min and the cell pellet rinsed with 1 x phosphate-buffered saline (PBS) before storage. Semen samples were split into 50-100 µl volume aliquots which were rinsed with 1 x PBS (5000 x *g* for 5 min). Mouth swabs were kept frozen until extraction, when they were resuspended into 100 µl PBS.

Genomic DNA was extracted from all the collected family samples (2 saliva lysates, 2 whole blood samples, 2 urine cell pellets, 6 buccal swabs, 1 semen lysate) on the Maxwell RSC Instrument using the Maxwell RSC Blood DNA kit (both Promega, WI, USA) and following manufacturer’s protocols. Aliquots of semen were pre-incubated in sperm lysis buffer (20 mM Tris HCl pH 8.0, 20 mM EDTA, 200 mM NaCl, 1% SDS) in the presence of proteinase K (250 μg/ml), dithiothreitol (DTT; 100 mM) and 0.6% SDS at 42°C for 4-12 hours. Concentrations of the final DNA eluates were assessed with standard fluorometric methods.

### Genotyping assay for verification of familial relationship using molecular inversion probes (smMIP assay)

To confirm the familial relationships of each trio, we used an in-house custom single-molecule molecular inversion probes (smMIPs) genotyping assay to capture common single nucleotide polymorphisms (SNPs) across all chromosomes (total of 290 smMIP probes targeting 154 autosomal, 14 X-linked and 57 Y-linked markers; for SNP details and probe sequences, see Supplementary Tables S4A & S4B) following established smMIPs protocols^2^ followed by sample barcoding, library preparation and 2 × 151 bp paired-end sequencing on a MiSeq instrument (Illumina, CA, USA). For each family, DNA from the proband sample obtained from the original diagnostic laboratory (or if unavailable, buccal swab DNA), the maternal blood sample and the paternal semen sample were analyzed. Sequencing data was processed using the ‘pileups snps’ tool in the amplimap v0.4.9^3^ pipeline with default settings (alignment to GRCh38.p12 with BWA, variant calling with GATK) to generate counts for the reference (REF) and alternate (ALT) alleles at each locus. Subsequently, the autosomal and linked SNP genotype for each individual of the family trio was recorded as Homozygous REF (AA), Heterozygous (AB) or Homozygous ALT (BB). For genotyping, SNPs were considered informative when the parents were homozygous (AA or BB) and the proband exhibited the expected genotype such as when Parent1/Parent2/Proband were AA/AA/AA, BB/BB/BB, AA/BB/AB. Other SNPs were analyzed to ensure there was no genotype discordance across the 3 family members.

### Ultra-deep Illumina sequencing (Deep-NGS) of DNM sites

Ultra-deep Illumina sequencing was performed in order to detect low levels of mosaicism in parental samples or post-zygotic mosaicism in the child. For each family-specific DNM, a pair of PCR primers tailed with generic CS1 (5’- ACACTGACGACATGGTTCTACA) and CS2 (5’- TACGGTAGCAGAGACTTGGTCT) sequence tags was designed to amplify a short genomic region (49-266 bp) around the DNM site; primer genomic locations (build GRCh38.p12), are provided in Supplementary Table S1. Each primer set was tested on control DNA with either High Fidelity Phusion or Q5 Polymerase (New England Biolabs, MA, USA) and PCR amplification was performed following manufacturer’s recommendations using 30 ng of genomic DNA from triplicates of up to 14 biological samples and three unrelated control DNAs in 10 µl PCR reactions, applying an initial denaturation step for 30 s at 98 °C, followed by 30 cycles of 10 s at 98 °C, 30 s at 68 °C, and 30 s at 72 °C, and 8 min at 72 °C as final extension step. Successful amplification was confirmed by running samples on an agarose gel. PCR-amplified fragments were diluted, further PCR amplified using individually barcoded primers, pooled together to construct libraries and ultra-deep sequenced, as previously described^4^ on a MiSeq (Illumina) instrument with 2 × 151 bp paired-end reads at an average depth of ∼19,000 x for each sample.

### Deep-NGS data analysis and determination of the observed variant allele frequency (VAF) at the DNM location

Illumina data were analyzed using amplimap^3^, as above, to obtain both the variant allele frequency (VAF) of each family-specific mutation and the total count of >Q30 bases at the corresponding genomic position (GRCh38.p12) in each PCR replicate and sample. For each family-specific dataset, DNM VAFs observed in each sample were corrected, to account for the background alternate read counts observed in the control samples (false-positives) at the DNM genomic location. Let *k1* and *k2* be the number of alternate reads observed in the control and case, and *n1* and *n2* be the total number of reads observed in the control and case, respectively. Let *p* denote the unobserved proportion of cells carrying a variant and let *q* be the false-positive rate of the sequencing and variant-calling procedure.

The joint likelihood of *p* and *q* is defined as follows

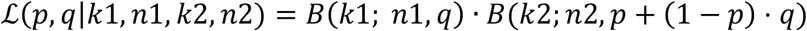

where *B* denotes the binomial probability mass function and *B*(*k; n, p*) is the probability of observing *k* successes in *n* trials with success probability *p*. The first term corresponds to the probability mass of observing k1 false-positives in the control, and the second term corresponds to the probability mass of observing *k2* alternate reads in the *p* + (1 − *p*) · q case. The rate in the second term corresponds to the fact that a read identified as carrying the variant in the case is either a true positive (i.e. actually carrying the variant) with probability *p* or a false positive (i.e. background noise, not carrying the variant but mistakenly identified as doing so) with probability (1 − *p*) · *q*).

We treated *q* as a nuisance parameter and obtained the marginal likelihood of *p* by numerically integrating the joint likelihood over *q* using adaptive quadrature^5^. Finally, we obtained the maximum likelihood estimate of *p* by numerically maximizing the marginal likelihood and obtained 95% confidence intervals using profile likelihood^6^. Scripts describing this analysis are available at github.com/sjbush/pregcare.

### Allele-specific PCRs

For two DNMs – a 44 bp deletion in *MECP2* in FAM12 and a 35 bp duplication in *MAGEL2* in FAM54 – the regions were successfully amplified as described above, but the deep-sequencing on the MiSeq platform did not lead to quantifiable results in the proband sample, making the assay unsuitable for mosaicism detection. Therefore, individual mutation-specific PCR assays were designed and the resulting PCR products analyzed using gel electrophoresis. The individual assays’ sensitivity was determined with dilution series experiments (Supplementary Note 2). Furthermore, an allele-specific PCR had to be designed for haplotyping the DNM of FAM38 in *AHDC1* due to a homopolymeric region around the mutation site for which the mutant and wildtype allele could not be phased with ONT sequencing (Supplementary Note 3 and Supplementary Table S3B).

### Long-read haplotyping assay using Oxford Nanopore Technologies (ONT)

The MinION (Oxford Nanopore Technologies [ONT], UK) long-read sequencing technology was used to determine the parent-of-origin of the DNM in the proband. To do so, primers were designed to amplify two regions (∼2-16 kb each, for locations of individual primer sequences, see Supplementary Table S1) on either side of the DNM. DNA from the two parental blood samples and the diagnostic genomic DNA from the proband were amplified using LongAmp Polymerase (New England Biolabs, UK) starting with 50 ng genomic DNA in a 20 µl reaction following manufacturer’s recommendations and the cycling conditions: initial 2 min at 95 °C, 30 cycles of 30 s at 95 °C and 16 min at 65 °C, and a final extension at 65 °C for 20 min. PCR amplicons were checked on a 0.9% agarose gel and if amplification had been successful, regions 1 and 2 from one sample were pooled. For library preparation, the PCR barcoding amplicon protocol and 1D ligation kit and PCR expansion kit (all ONT, UK) were used to barcode individual samples in a 20 µl PCR reaction with LongAmp polymerase, 2 µM barcoding primers and 1:100 diluted target PCR with the cycling conditions as described above for 8 cycles. After adapter ligation, the pooled library was loaded onto a MinION SpotOn Mk I version R9 flowcell (ONT) for sequencing following the manufacturer’s recommendations. For initial data processing (demultiplexing and basecalling) each set of fast5 files was processed using Guppy v4.5.4+66c1a77 (https://community.nanoporetech.com) with the parameter --config dna_r9.4.1_450bps_hac.cfg, producing one set of reads for each barcode/family member of the trio. Reads are deposited in the European Nucleotide Archive under BioProject accession number PRJEB53977 (http://www.ebi.ac.uk/ena/data/view/PRJEB53977).

### Haplotype phasing of *de novo* mutations using Medaka and mpileup

ONT reads for each trio were aligned to the GRCh38.p12 primary assembly using minimap2 v2.18^7^ with parameter -ax map-ont. Lower-quality (MAPQ < 20) and non-primary alignments were discarded using samtools view v1.12^8^ with parameters -q 20 -F 256 -F 2048. For each target region (genomic coordinates are given in Supplementary Table S1), variants were called using the ‘medaka_variant’ workflow of Medaka v1.3.2 (https://github.com/nanoporetech/medaka, accessed 6th May 2021) with default parameters. The set of VCFs per region were then concatenated using BCFtools v1.12^8^ to produce one VCF per BAM, subsequently annotated using dbSNP v153^9^ (https://ftp.ncbi.nih.gov/snp/latest_release/VCF/GCF_000001405.38.gz, accessed 6th May 2021).

Where possible, Medaka uses the information contained within heterozygous SNPs to impute the haplotype of the aligned reads. In practice this means that a proportion of the calls in each VCF are phased, being assigned to a ‘phase set’ of SNPs on the same haplotype. Given that the sequencing data represent mother/father/proband trios, with each proband having a DNM, each VCF was parsed to determine whether Medaka had called and phased the DNM in the proband (but not in either parent, confirming their true “de novo” status). For each DNM called by Medaka, we obtained the associated phased set SNPs, retaining only those which had a total depth of coverage >10x. We cross-referenced the phased set SNPs with the VCFs from the mother and father and identified which calls (if any) had been made at those positions. This produced a set of three haplotypes from which we used a custom script to classify the inheritance of the DNM as either maternal or paternal (the SNPs in phase with the DNM could only be derived from the chromosome inherited from the mother or father, respectively), else unresolved (Medaka either did not call the DNM in the child, called it but did not construct a phased set, or, if it did construct a phased set, either did not call its constituent SNPs in the parents or made identical calls for both of them).

DNMs not successfully phased using Medaka (Supplementary Note 3) were phased by programmatic and/or manual inspection of read pileups. A programmatic approach was implemented using a custom script which parsed read pileups (generated using samtools mpileup with parameters -aa --output-QNAME) to obtain a set of reads which contained both the ALT-allele for the DNM and a candidate phasing SNP (considered the closest one to it and for which there was a prior, namely inclusion in dbSNP). We then constructed a 2×2 count table (rows: number of reads calling REF/ALT at DNM position, columns: number of reads calling REF/ALT at phasing SNP positions) and resolved inheritance by identifying which of the two alleles for the phasing SNP, REF or ALT, were disproportionately found on the same read as the DNM ALT. Significance was assessed using Fisher’s exact test.

Haplotypes flagged as not programmatically resolved by either Medaka or pileup were manually reviewed using IGV v2.11.2^10^, with visual inspection also used to validate all the above calls. Full details, and all scripts used for this analysis, are available at github.com/sjbush/pregcare. For one family (FAM38) the phase could not be resolved with long read-sequencing due to a homopolymeric stretch around the mutation site. For this family, an allele-specific PCR was performed (Supplementary Note 4).

