## Supplementary Figures and Notes for "The PREGCARE study: Personalized recurrence risk assessment following the birth of a child with a pathogenic *de novo* mutation"

### List of Supplementary Material

**Supplementary Figures**

Supplementary Fig. S1: The timing of occurrence of the DNM impacts on recurrence risk.

Supplementary Fig. S2: Deep-sequencing results for all biological samples analyzed in the mosaic families.

**Supplementary Tables**

Supplementary Table S1. Overview and characteristics of the 61 DNMs from the families enrolled in the PREGCARE study

Supplementary Table S2. Deep-Sequencing (Deep-NGS) data for the families enrolled in the PREGCARE study

Supplementary Table S3A. Overview of the phasing strategy for each of the individual PREGCARE families

Supplementary Table S3B. Description of the informative SNP and read count tables supporting the phasing of DNMs and parent-of-origin determination for individual PREGCARE families

Supplementary Table S4A. Details of the common single nucleotide polymorphisms (SNPs) used for genotyping the members of family trios and verifying the family relationship (smMIP assay)

Supplementary Table S4B. Sequence information of the Molecular Inversion Probes (smMIP assay)

**Supplementary Notes**

Supplementary Note 1: Stratification of DNMs into 7 categories

Supplementary Note 2: Design and results for assessment of mosaicism of two families with the larger de novo indels (FAM12 and FAM54)

Supplementary Note 3: Successes and Failures to resolve haplotype with ONT sequencing

Supplementary Note 4: Allele-specific PCR for haplotyping the DNM in *AHDC1* in FAM38

Supplementary Note 5: Estimating the recurrence risk associated with mosaicism from multi-sibling families and sperm WGS data.

<u>**Supplementary Figures**</u>

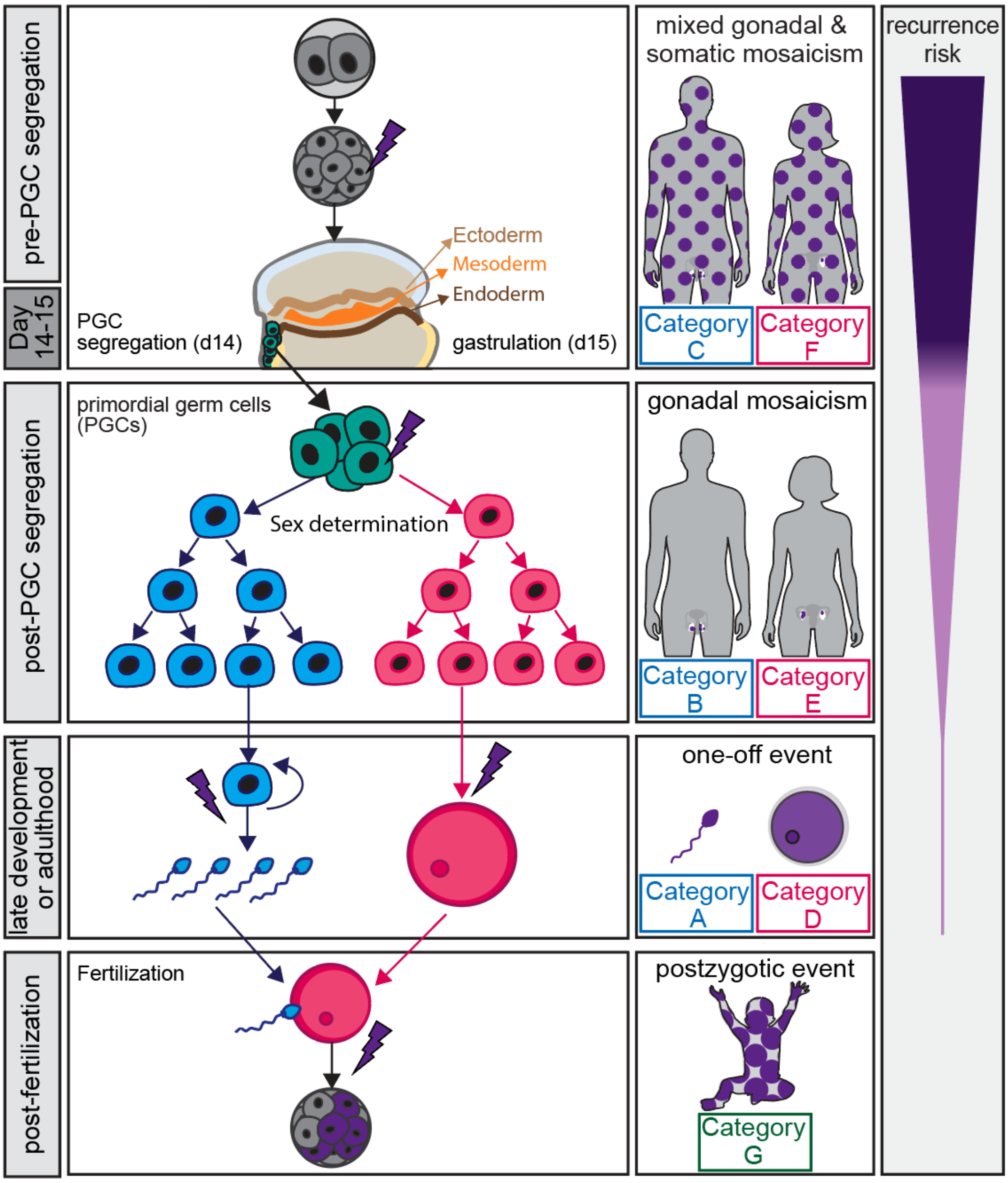


**Supplementary Fig. S1: The timing of occurrence of the DNM impacts on recurrence risk.** *De novo mutations (DNMs) can occur at any point prior to, or during the development of the embryo, potentially resulting in mosaicism. If the DNM arises in a parent prior to the segregation of primordial germ cells (PGCs) and gastrulation (days 14-15 of embryogenesis), it can result in mixed mosaicism affecting both the somatic tissues and gonads in either the father (Category C in this study) or the mother (Category F); because they occur very early, these scenarios are associated with the highest sibling recurrence risk. If the DNM arises in parental PGCs after PGC segregation, mosaicism will be confined to the gonads in the father (Category B) or the mother (Category E); circumstances associated with intermediate sibling recurrence risks, depending on the relative proportion of mutant cells. Because mosaic events occur before sex determination, they affect both genders equally. If the DNM arises as a one-off mutational event in late development or adulthood in the father (Category A) or the mother (Category D), the recurrence risk is negligible. The DNM may also arise as a post-zygotic event in the child following fertilization (Category G); in this case, the recurrence risk is zero, as the DNM is not present in the parental gametes. It is possible to quantify the recurrence risk for DNMs in Categories A-C and G, but the recurrence risk cannot be quantified for DNMs of maternal origin (Categories D-F) as oocytes cannot be accessed without invasive sample collection.*

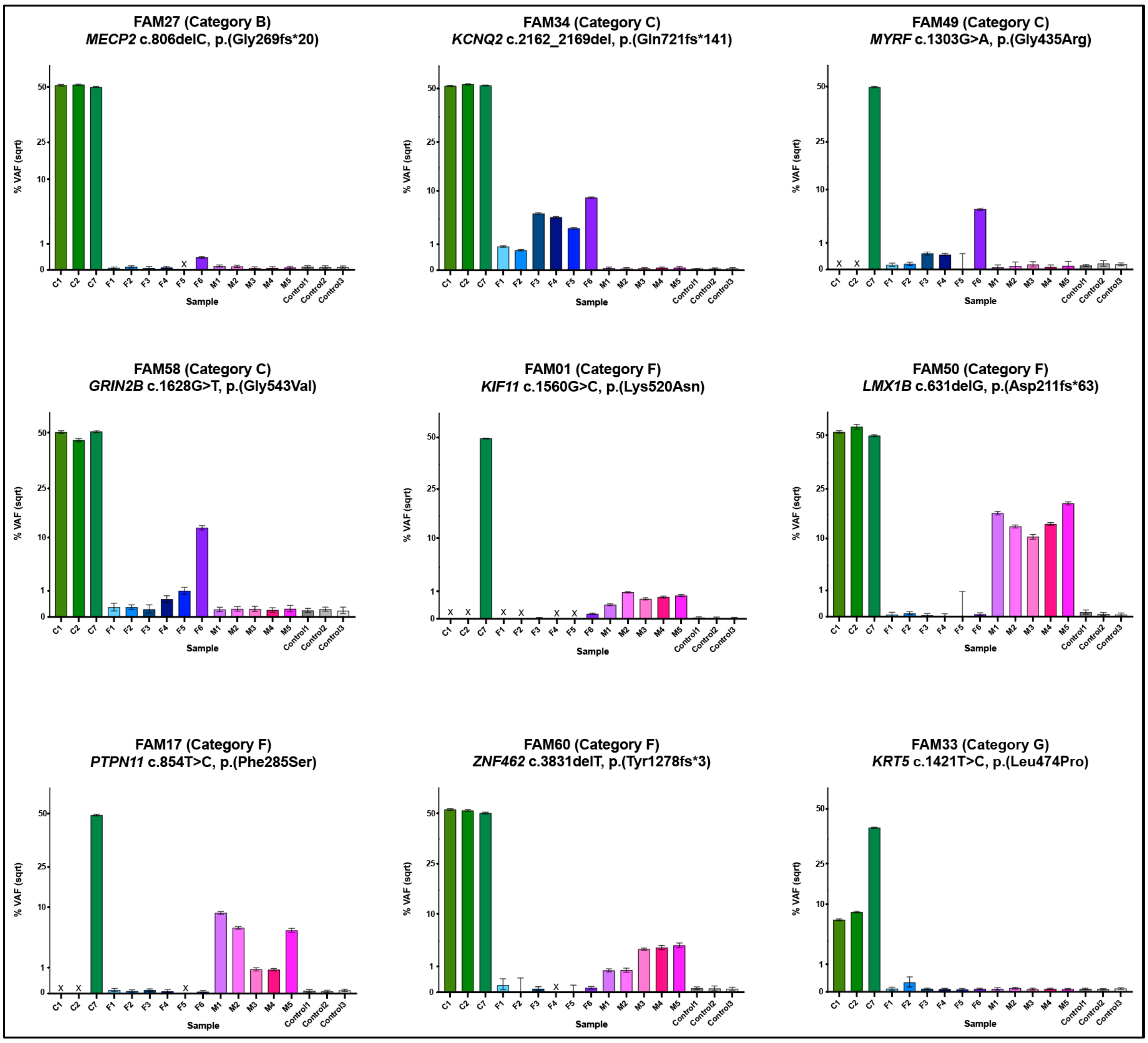


**Supplementary Fig. S2: Deep-sequencing results for all biological samples analyzed in the mosaic families**. *Variant allele frequencies (VAF) measured for each tissue sample analyzed from the three family members and three unrelated controls plotted as the square root (sqrt). Family number, Category, affected gene and coordinates of the DNM are indicated above each plot. The origin of the different tissue samples is indicated on the x-axis and colors have been used for ease of visualization with samples from C=child (green); F=father (shades of blue/purple); M=mother (shades of pink/red); Control (grey). VAFs are average percentage of mutant reads measured by deep-NGS and error bars represent the 95% binomial confidence intervals across three technical replicates (all data are provided in Supplementary Table S2). Abbreviations used for sample identification: 1=buccal swab (left); 2=buccal swab (right); 3=blood; 4=saliva; 5=urine; 6=sperm; 7=gDNA from original testing; X represents a missing or failed sample.*

### PREGCARE: Supplementary Notes

The PREGCARE study: Personalized recurrence risk assessment following the birth of a child with a pathogenic de novo mutation

Bernkopf, Abdullah *et al.*

**Content:**

Page 2: Supplementary Note 1: Stratification of DNMs into 7 categories

Page 5: Supplementary Note 2: Design and results for assessment of mosaicism of two families with the larger de novo indels (FAM12b and FAM54)

Page 7: Supplementary Note 3: Successes and failures to resolve haplotype phasing using ONT sequencing

Page 9: Supplementary Note 4: Allele-specific PCR for haplotyping the DNM in *AHDC1* in FAM38

Page 10: Supplementary Note 5: Estimating the recurrence risk associated with mosaicism from multi-sibling families and sperm WGS data

Page 15: References

**Supplementary Note 1: Stratification of DNMs into 7 categories**

To determine the relative proportion of cases in the seven categories of *de novo* mutation (DNM) origin presented in Figure 1, we used the data described in Rahbari *et al* (2016)[1]. In this study, a total of 768 DNMs were identified across whole-genome sequencing (WGS) data (average depth: 25x) for three multi-sibling families comprising 4-5 individuals and their parents. Among these, 399 DNMs could be haplotyped in respect to a nearby inherited polymorphism and allowed the parent-of-origin to be determined, demonstrating that 78% (311/399) of DNMs were paternal in origin, a result consistent with other studies[2-5].

The proportion of DNMs that were likely to have originated through parental mosaicism could be determined via two different approaches:

(a) If mosaic mutations are present in multiple parental gametes, recurrence in siblings may be observed. In the Rahbari study[1], ten validated DNMs were shared by at least two siblings of the same family, providing an estimate of any germline mutation being shared by at least two siblings as 1.3% (10/768). However, some of the mosaic variants were shared by more than two siblings in the family. Hence, another, more exact, way to estimate the observed recurrence of DNMs in these 3 families, proposed by Rahbari *et al.* (Supplementary Table S2), described a total of 2850 sib-sib comparisons (sum of number of DNMs multiplied by the number of other siblings in that family, for each given DNM) and 34 instances of shared DNMs between sib pairs, leading to a figure of 1.19%, the probability that a given DNM is also present in another sibling (i.e. the recurrence risk for a DNM that has already been observed in one individual of the family, conditional on parental blood VAFs being not detected at a depth of ~25x, a sequencing depth estimated to have a sensitivity ~10%). Overall, this genome-wide estimate of DNM recurrence, although based on a small number of multi-sibling families, provides a figure in line with the empirical 1-2% population recurrence risk[6,7].

(b) A mosaic (parental) origin for validated DNMs was also sought by deep-sequencing (average 567x) of parental blood DNA (note, blood was the only tissue available for this study)[1]. This approach showed that 25 DNMs observed in children were in fact detectable at low VAF (observed average VAF of 3.2% (range: 0.6-10.2%)). Of note, the pipeline used for calling the 768 DNMs in the three families conservative balanced the need for sensitivity and specificity in this dataset, as demonstrated by the low VAFs observed by deep-sequencing for the 25 mosaic mutations in parents which were all ≤10.2%. After correction for incomplete power to detect parental mosaics at VAF < 1%, these data suggest that ~4.2% of DNMs observed in children are present in the somatic tissues (mixed mosaicism) of one of the parents. As expected for early mutational events, these were approximately equally distributed among the originating parent (paternal:maternal ratio = 9:16), a result consistent with other estimates[4]. This implies that, overall, paternal (Category C) and maternal (Category F) mixed mosaicism are each responsible for ~2% of the DNMs occurring in a child (Fig. 1).

By comparing the two approaches above, the contribution of confined gonadal mosaicism (Categories B and E) can be deduced, as among the ten recurrent DNMs, six could also be detected in the blood of one of the parents (i.e. mixed mosaic) and four were undetectable at ~500x (i.e. indicative of confined gonadal mosaicism). From this, it can be inferred that the proportion of mixed and gonadal mosaic mutations are likely comparable and corresponds to ~4% each, split equally between the two parents, meaning that each of the Categories B, C, E and F corresponds to ~2% of DNMs (Fig. 1).

The proportion of DNMs belonging to Category G (high-level VAF post-zygotic mutation having occurred after fertilization in the proband) is based on data from Acuna-Hidalgo *et al.* (2015)[8] who analysed a set of 107 DNMs from 50 parent-offspring trios, using four different sequencing techniques, and showed that seven (6.5%) of these apparent DNMs had VAF significantly different to the 50:50 ratio expected for heterozygous germline variants. These post-zygotic Category G variants are expected to arise equally on the maternal and paternally derived chromosomes.

Taking all these data into consideration allows us to estimate the representation of the different mosaic events into each category A-G as presented on Figure 1. The contribution of Categories A and D (one-off events of paternal or maternal origin, respectively) can be deduced, given that globally 78% of DNMs are paternal (and 22% are maternal) in origin. Overall, these data suggest that ~8% of all candidate DNMs identified in a proband are in fact present as mosaic in the parental germline (Categories B-C or E-F) and represent a recurrence risk in a future pregnancy. Of note, given that 78% of DNMs are paternal in origin this also implies that a mosaic origin is expected 3-4x more frequently for a DNM of maternal origin than for paternally-derived DNM.

Although the genome-wide estimates of the occurrence of different categories of DNM used in Figure 1 are based on studies[1,6] of relatively small sizes, these estimates are concordant with more recent data[4,5]. The 33 large (~8 siblings) three-generation families analyzed in the Sasani *et al.* study[4] allowed them to show that 1.3% (303/23,386) of DNM sites are shared by one or more (up to 5) siblings of the same family. To refine this estimate, we used the Sasani dataset and performed the same analysis as that described in Supplementary Table S2 of Rahbari[1] (see above) to establish the proportion of a DNM being shared amongst any two siblings (conditioned on the fact that the DNM was not detected in parental blood at ~30x sequencing depth). We considered the 23,386 DNMs called in the multi-sibling third generation and the observed 720 shared mosaic events (across 303 unique sites). These represent a total of 1156 sharing events (sum across individuals of number of shared occurrences of a DNM [i.e. for each shared mutation, count how many individuals (other than themselves in this family) carried this DNM]) from a total 221,569 sib-sib comparisons (sum across individuals of number of sibs [not including themselves] * [number of DNMs]), which provides an observed recurrence between any two siblings of 0.52%, a factor ~2 lower than that from the analysis of the three families described in Rahbari[1]. Of note, the 1.19% average estimate from the Rahbari study[1] was mainly contributed by one of the three families analysed with individual values of 0% (0/780), 0.3% (2/706) and 2.3% (32/1364).

Finally, it is worth mentioning that the unique three-generation design of the Sasani study[4] also provides an estimate of the proportion of high-level post-zygotic mosaic cases (Cat G) caused by very early events (before the soma-germline split) in the second-parental generation. These mosaic DNMs will exhibit high VAFs that are difficult to distinguish from the 50:50 true heterozygous germline DNM (such as those of Category G in our study). By looking at the transmitted haplotype around the DNM called in the second-parental generation within the third-generation probands, the authors can show that 9.3% of DNMs (475/5017 DNMs) observed in the second generation are, in fact, post-zygotic mosaic in these individuals.  This also allows them to confirm that, as expected, this process is sex-independent as each parent contributes equally to this tally (paternal:maternal ratio = 249:226).

**Supplementary Note 2: Design and results for assessment of mosaicism in two families with the larger indel DNMs (FAM12 and FAM54)**

For FAM12, one of the proband's DNM was a 44 bp deletion in *MECP2* (FAM12b) for which we designed a mutation-specific PCR assay using a reverse primer encompassing the deletion junction with the 5' sequence starting at g.X:154,030,684 and the sequence 5'-CTGCTCCCACCCCTGCCtCa**CTG**-3', where t and a represent introduced mismatches and the sequence in bold (CTG) represents g.X:154,030,618_154,030,620, located on the other side of the deletion. PCR in combination with a forward primer (g.X:154,030,385-154,030,404) (Tm = 66º C) was able to amplify specifically the mutant fragment containing the 44 bp deletion and the presence of the mutant band (256 bp) was assessed on a 3% agarose gel. Serial dilution of the proband gDNA to levels down to 1:650 (using carrier DNA), showed robust amplification of the mutation-specific fragment (256 bp) at this dilution, demonstrating assay sensitivity to levels below 0.08% (1:1,300). In parallel, a control PCR flanking the deletion (using the following primers: FW1: g.X:154,030,578-154,030,597 and Rev1: g.X:154,030,796-154,030,777) was performed to ensure robust amplification of all the family samples tested and to assess the presence of the two alleles (generating fragments of 155 bp (mutant) vs. 199 bp (WT)) in the expected ratio in the proband's samples. Consequently, there is no evidence suggestive of a post-zygotic mosaic event in the proband. Using the mutation-specific PCR assay, a robust amplification of the 256 bp band was observed for all the three proband samples (left and right buccal swabs and blood), but no amplification of the parental samples (blood, saliva, buccal swabs, urine and semen) was detected; therefore, no evidence of mosaicism was detected in any of the parental samples.

For FAM54 the DNM was a 35 bp duplication in *MAGEL2* and two PCR assays were used to screen all the family samples and assess the results on agarose gels. First, a genomic region flanking the duplication (using the forward primer g.15:23,645,351_23,645,372 and reverse primer 1: g.15:23,645,608_23,645,585) was defined to ensure robust amplification of all the family samples tested and to assess the presence of the two alleles. By inspection, the two fragments (293 bp (mutant) vs. 258 bp (WT)) were present in the expected ratio in the proband (no evidence to suggest post-zygotic mosaicism), while a single band of 258 bp (and no mutant 293 bp) was observed in the parental or control samples. We then designed an allele-specific PCR assay (using a reverse primer encompassing the duplication junction with 5' sequence starting at g.15:23,645,498 (5'-CGCATGATCTTTGCTGC**AGG**-3', where the sequence in bold (AGG) represents g.15:23,645,516-23,645,514, located specifically within the duplication) in combination with the forward primer as described above, able to amplify specifically the mutant fragment containing the 35 bp duplication. The presence/absence of the mutant band (183 bp) was assessed for all the family samples on a 3% agarose gel. Moreover, serial dilution of the proband gDNA to levels down to 1:650 (using carrier DNA), showed robust amplification of the mutation-specific fragment (183 bp) at this dilution, demonstrating assay sensitivity to mutation levels below 0.08% (1:1,300). Using the mutation-specific PCR assay, robust amplification of the 183 bp band was observed for all the three proband samples (blood, left and right mouth swabs), but no amplification of the parental samples (blood, saliva, mouth swabs, urine or paternal semen) or in control samples was detected; therefore, we concluded that no evidence of mosaicism was found in any of the parental samples.

**Supplementary Note 3: Successes and failures to resolve haplotype phasing using ONT sequencing**

In total, 50 family trios were sequenced with ONT. These included 47 families without evidence of mosaicism as presented in the main text and three families in which we detected mosaicism and were included as controls (FAM17 (Cat F), FAM27 (Cat B) and FAM34 (Cat C)). In this series, DNMs consisted of 37 SNVs and 13 indels. Overall, we were able to resolve the phase for 41 DNMs.

The workflow used to haplotype DNMs is available at github.com/sjbush/pregcare and sequentially implements the programs Medaka and mpileup. In brief, if Medaka fails to resolve phase, the workflow will attempt to do so using pileup. Should phase not be resolved using pileup, manual phasing is attempted.

More specifically, Medaka resolved the inheritance pattern for 21 of the SNVs (56.8% of the total SNVs) but none of the indels. Poorer performance with indels was likely due to the relatively error-prone nature of ONT sequencing data[9]. Furthermore, according to ref.[10], Medaka first predicts SNVs from unphased reads and then uses WhatsHap[11] to phase them. Indel calling – technically more challenging – was performed later, and only using reads already phased, a subset of the total. Through data curation, failures to resolve the inheritance of the SNVs with Medaka (16/37) could be attributed to the DNM not being called in the child (six families), the DNM being called but not assigned a phase set (five families), and to the set of in-phase SNPs being either identical in both parents (four families) or considered by Medaka to be low-quality calls (one family). Failures to resolve the inheritance of indels with Medaka could be attributed to the DNM not being called in the child (12 families) and the DNM being called but not assigned a phase set (one family). We found no substantive improvement in the number of DNMs phased when substituting Medaka for the alternative variant callers of Clair3 v0.1-r7 (https://github.com/HKU-BAL/Clair3) or PEPPER-Margin-DeepVariant v0.5[12] (data not shown). Of the 29 families for which inheritance was not resolved using Medaka, we resolved inheritance using pileup in 15 cases (see Methods and Supplementary Table S3A & S3B). For one family (FAM02), the informative SNP was an indel and the DNM was phased manually. For a further three families (FAM11, FAM38 and FAM67), the phase was resolved manually using a ‘2-SNP’ design; i.e. we first identified a SNP that was heterozygous in all three family members and used another discriminant SNP heterozygous in one of the parents to assign the phase of each allele to their parent-of-origin (detailed in Supplementary Table S3B).

For the nine remaining families (5 SNVs and 4 indels), we were unable to resolve the phase of the DNM. A key requirement for haplotype phasing of DNMs is the presence of a heterozygous SNP in the vicinity of the DNM in the proband, in order to distinguish the two parental alleles. In all nine cases, no heterozygous SNP could be identified near the DNM in the proband, despite sequencing a total of >10 kb around the DNM (and >20 kb in 7 families; see Supplementary Table S3A). In theory, these DNMs can be haplotyped, but this will require further extension of the size of the genomic region interrogated beyond the continuous runs of homozygosity. Implementation of ultra long-read technologies, such as those afforded by the Telomere-to-Telomere (T2T) sequencing approach, will undoubtedly improve our ability to phase DNMs in the future. In a recent study, Noyes *et al.*[13] showed that 194/195 DNMs identified in a family quartet could be assigned to their parent-of-origin, using informative SNPs extending 20 kb on either site of the DNM location.

#### Supplementary Note 4: Allele-specific PCR for haplotyping the DNM in *AHDC1* in FAM38

For FAM38, the ONT sequencing did not allow unambiguous discrimination of the two alleles at the DNM position g.1:27548981 (deletion G) in the *AHDC1* gene in the proband due to a homopolymeric region (GGGG (mutant) vs. GGGGG (REF)). Using the ONT data, we identified a SNP (rs2076457) that was heterozygous in the three members of the family. We could establish the parent-of-origin of each of the two rs2076457 alleles (paternal A and maternal C) in the proband by using the phase in respect to another discriminant SNP (rs113173951G/A) which was heterozygous in the mother only. Hence, the rs2076457 SNP was used for haplotyping the *de novo* mutation by performing an allele-specific PCR amplification followed by dideoxy-sequencing. To amplify only the maternally-derived rs2076457C allele in the proband, the reverse primer 5′-CAGCCGCTGGGGTCGGGGGCaCG-3′, where a represents a deliberate mismatch and G is rs2076457C allele-specific, was combined with a forward primer (g.1:27,548,900_27,548,920), to generate a fragment of ~1.1 kb. Dideoxy-sequencing of this PCR fragment revealed that the maternally-derived rs2076457C allele is in phase with the *de novo* mutation (g.1:27,548,981delG) (see Supplementary Table S3B).

**Supplementary Note 5: Estimating the recurrence risk associated with mosaicism from multi-sibling families and sperm WGS data.**

In what follows, we define recurrence risk (RR) to be the probability that a given DNM observed in a proband is present in the next offspring from the same couple. Notably, this means the estimates of recurrence risk obtained below would be inappropriate if sequencing results from another offspring were known. We further condition our estimates on the fact that the DNM has been called in the proband (i.e. HET) and was not called (i.e. HOM REF) from both parents, given that they had been sequenced using NGS at ~25-30x depth. Importantly, there is relatively limited sensitivity of variant calling using routine NGS, where calling a mosaic VAF = 15% would require detection, and filtering in, of only 3-4 mutant reads in a parent sequenced at 25x[14]; this sensitivity is similar to the 15-20% limit of detection of dideoxy-sequencing [15].

We used the estimates of the proportion of DNMs belonging to each category A through G, as described on Fig. 1 and in Supplementary Note 1.

Next, we used data from Yang *et al.*[16] that describes deep (200-300x) Whole Genome Sequences (WGS) of paired blood and sperm samples from 17 men, to derive estimates of the distribution of gonadal variant allele frequencies (VAFs) for mosaic variants, conditional on them being either detected in both sperm and blood (i.e., mixed mosaic corresponding to Category (Cat) C and by inference Cat F), or just being mosaic in sperm (i.e., confined gonadal mosaic corresponding to Cat B and by inference Cat E). Of note, in the absence of direct estimates of female gonadal mosaic frequencies, given that these are very early events, we assume that the distributions of VAFs are the same for Categories B and E, and likewise for Categories C and F (see also main text and Supplementary Fig. S1).

We downloaded Data S1 of Yang et *al.*[16], and for gonadal tissue only (Categories B (and E)), used entries from COHORT = "Young Age" or "Advanced Age"; SET_SPERM_ONLY = 1, and used column MAF_SPERM_A. For mixed mosaic (Categories C (and F)), we used entries from COHORT = "Young Age" or "Advanced Age"; SET_BOTH_MOSAIC = 1, and again used column MAF_SPERM_A. We further stratify the mosaic data by restricting them to MAF_BLOOD < 0.15 as an approximation, as variants with higher mosaic VAFs in blood are less likely to be assigned as DNMs (for which the parental sample must be called as HOM REF) in WGS at ~25-30x, which would violate the earlier assumption. We also perform the calculations without this MAF_BLOOD < 0.15 restriction, to model an upper bound of recurrence when blood is not used for WGS and/or VAFs are very variable in different somatic tissues. Notably, we did not include the data from COHORT = “ASD” as these samples were sequenced at a lower depth (200x vs. 300x)[16].

We obtained an estimate of the average VAF for gonadal (Cat B and Cat E) and mixed mosaic (Cat C and Cat F) as follows:
For SPERM-only variants (n = 418): $VAF_{Sperm}$ = 3.01%
For MIXED mosaic (all variants) (n = 190): $VAF_{Sperm}$ = 8.39%
For MIXED mosaic (for variants with $VAF_{Blood}$ <0.15) (n = 151): $VAF_{Sperm}$ = 4.89%

To calculate the recurrence risk, we make the assumption that each DNM in a given category must be of a given “type”, and that each DNM of the same type has the same underlying VAF in the originating parent’s gonadal tissue. We model each observation in the Yang data as akin to a DNM type, with $VAF_{sperm}$ as the probability of observing the non-reference (ALT) allele for that DNM type. Using Bayes Theorem, we model that the probability a DNM is type i (i.e. when observed in a given category), as being proportional to the VAF of a DNM of type i from that category. Specifically, here we model that the probability of an observed DNM of a given category is type i (i in 1 through the length of the Yang $VAF_{sperm}$ data for that category) is the frequency of that type as specified in the Yang data, divided by the total frequency for that category (as given in the Yang data).
We therefore calculate the recurrence risk (RR) as follows: where R is the event of recurrence (i.e. a ALT allele for the observed DNM is present in the next offspring at the same genomic site); DNM obs is the event of observing a DNM at an otherwise arbitrary site. Then we have that: P(Cat = X | DNM obs) is the previously described probability of the observed DNM belonging to categories A through F; P(Type = i | Cat = X, DNM obs) is the previously described probability that an observed DNM of category X has the property of type i (i.e. has true underlying gonadal allele frequency of type i); and P(R | Type = i, Cat = X, DNM obs) is the probability of recurrence given an observed DNM of category X and type i, which has gonadal allele frequency of type i.

$$RR = P(R \mid DNM\ obs)$$
$$= \sum_{X\ in\ A:F} P(R, Cat = X \mid DNM\ obs)$$
$$= \sum_{X\ in\ A:F} P(R \mid Cat = X, DNM\ obs)\, P(Cat = X \mid DNM\ obs)$$

$$= \sum_{X\ in\ A:F} \sum_{type\ i} P(R, Type\ =\ i \mid Cat\ =\ X, DNM\ obs)\ P(Cat\ =\ X \mid DNM\ obs)$$

$$= \sum_{X\ in\ A:F} \sum_{type\ i} P(R \mid Type\ =\ i, Cat\ =\ X, DNM\ obs)\ P(Type\ =\ i \mid Cat\ =\ X, DNM\ obs) P(Cat\ =\ X \mid DNM\ obs)$$

Together, this allows us to estimate the recurrence risks as presented in the table below, for the overall population risk, as well as when the DNM belongs to a subset of categories (maternal origin without suspicion of somatic mosaicism (D/E)), unresolved parent-of-origin, without suspicion of mosaicism in parental tissues (A/D/E)) or mosaic cases (mixed C/F or gonadal B/E). Note, as explained above, we report results both when restricting Yang mosaic sperm variants with $VAF_{Blood}$ <0.15 which generally is likely to best represent the situation most families would fall into after routine clinical testing, as well as without this cut-off.

| **Category** | **RR % ($VAF_{Blood}$ <0.15)** | **RR % (all)** |
|---|---|---|
| Overall | 0.58 | 0.89 |
| D or E | 0.49 | 0.49 |
| A, D or E | 0.095 | 0.095 |
| C or F | 10.27 | 17.95 |
| B or E | 4.18 | 4.18 |

For comparison, we note that Rahbari *et al.*[1] estimated an overall RR = 1.2% in their Supplementary Table S2. By performing a similar calculation using substantially more data from Sasani *et al.*[4], we obtain an estimate of 0.52% (see Supplementary Note 1 for details), which is similar to our estimate of RR for $VAF_{sperm}$ <0.15 of 0.58% above.
From these data, it also follows that screening couples by deep-sequencing of multiple somatic tissues to detect the cases of parental mixed mosaicism (C or F) combined with sperm analysis to identify cases of paternal gonadal mosaicism (B), offer the possibility to reduce the remaining RR for the other couples (Cat A, D or E) to ~ 0.1%, representing approximately a 10-fold risk reduction from the starting overall population risk.

**Estimating confidence intervals on population risk estimates**

We estimated confidence intervals for the above estimates using bootstrapping. For each bootstrapping replicate, we re-sampled the variant allele frequency distributions from Yang *et al.*[16]. To form confidence intervals (CIs), we performed 100,000 bootstrap replicates, and took the 2.5th and 97.5th percentiles. This generated the following estimates and confidence intervals (in %)

| Category | RR % ($VAF_{Blood}$ <0.15) | RR % (all) |
|---|---|---|
| Overall | 0.58 (0.5, 0.64) | 0.88 (0.8, 0.96) |
| D or E | 0.49 (0.43, 0.57) | 0.49 (0.43, 0.57) |
| A, D or E | 0.09 (0.08, 0.11) | 0.09 (0.08, 0.11) |
| C or F | 10.23 (8.5, 11.81) | 17.91 (15.8, 19.83) |
| B or E | 4.16 (3.62, 4.82) | 4.16 (3.62, 4.82) |

We note that these CIs are likely insufficiently conservative (i.e. too narrow) as we are not correcting for intra *vs.* inter family variation in this bootstrapping, and we do not incorporate uncertainty in knowledge of the probability of the different categories.

We also note that the above are CIs, and as such, interval estimators that should contain the true underlying parameter with the appropriate confidence (here 95%). As new studies similar to the Yang *et al.*[16] dataset become available to better estimate CIs, these intervals would shrink to eventually reach a zero width. However, they do not reflect the fact that VAF varies across different families within the same category, resulting in significantly different levels of individual risk (i.e. when the VAF in gonadal tissue for a given DNM differ significantly from the sub-group average). We therefore calculated in each bootstrapping replicate from each of the sub-categories defined in the Table above, the recurrence risk corresponding to the 2.5th and 97.5th percentile of risk. We note that these estimates are not confidence intervals. Averaging these across bootstrapping runs yields the following (in %).

| Category | RR % ($VAF_{Blood}$ <0.15) | | RR % (all) | |
|---|---|---|---|---|
| | 2.5th ile | 97.5th ile | 2.5th ile | 97.5th ile |
| D or E | 0.0 | 5.0 | 0.00 | 5.0 |
| A, D or E | 0.0 | 0.0 | 0.00 | 0.00 |
| C or F | 1.0 | 20.7 | 1.5 | 33.5 |
| B or E | 1.4 | 13.5 | 1.4 | 13.5 |

Overall, we note that the RR estimates based on combining multi-sibling and sperm WGS data are similar but slightly lower than the 1-2% population risk quoted in the clinic. This generic figure is an empiric estimate, based on recurrence observed for different clinical disorders – often reported as case studies – that were ascertained using a variety of methodologies and for which it is difficult to find a reliable data source[6,7]. Moreover, depending on the disorders and associated mutated genes, the risk of mosaic presentation may be lower (i.e., paternal age-effect disorders, such as Apert syndrome or achondroplasia[17]) or higher (i.e., Duchenne and Becker Muscular Dystrophy[18], Osteogenesis Imperfecta[19], Dravet syndrome[20]) than the population risk for non-pathogenic mutations.
